# Influence of Monaural Auditory Stimulation Combined with Music on Brain Activity

**DOI:** 10.1101/2023.10.01.560062

**Authors:** Ming Chang, Kenta Tanaka, Yasushi Naruse, Yasuhiko Imamura, Shinya Fujii

**Affiliations:** Vie Style, Inc., Kanagawa, Japan

## Abstract

Recently, the increasing attention to mental states and psychophysical health has fueled the research into methods that can aid in relaxation and recovery. Traditional methods like meditation and sauna, while effective, have their limitations; thus, the need for more accessible and convenient alternatives. Our innovative approach combines monaural beats with music, attempting to replicate the relaxing effects of a sauna in the auditory domain. In comparison to normal music and silent condition, the power of the theta active band significantly increased when listening to our modified music. Furthermore, after listening to modified music, there was a significant increase in mismatch negativity (MMN) amplitude in the oddball task. Additionally, participants’ subjective responses to a questionnaire indicated significant changes in body relaxation and other metrics after listening to the processed music. This state is considered similar to the “totonou” state previously reported for the sauna. Thus, the present research proposes a convenient method for achieving relaxation, opening an avenue for individuals to customize their “totonou” music based on personal preferences.

## Introduction

In recent years, the growing emphasis on mental well-being and holistic health has driven the search for methods that facilitate relaxation and refresh. Meditation, an age-old practice for relaxation and mind and body attunement, is increasingly recognized in modern society, as meditation not only helps individuals achieving profound states of relaxation but also amplifies the theta or/and alpha brain activity [1-4]. However, its nuanced techniques and rigor can be challenging, leaving many struggling to genuinely immerse themselves in a meditative state.

Conversely, going to the sauna, an ancient practice for physical relaxation, is deeply ingrained in numerous cultures [5,6]. Compared to meditation, saunas do not require any specific techniques and can be enjoyed by anyone. Interestingly, recent studies indicate that saunas not only facilitate physical relaxation but also significantly increase alpha brain activity [7]. Moreover, our recent research [8] found increased theta and alpha waves after three sauna sessions. Moreover, in the oddball task conducted before and after, there was an increase in mismatch negativity (MMN) amplitude and a reduction in P300 amplitude after the sauna. This emphasizes that saunas can promote relaxation and enhance perceptual awareness, improving the overall sense of well-being, referred to as “totonou” state in Japanese culture. The effect of meditation and sauna in theta and alpha activity reveals the complex interplay between relaxation, cognitive rest, and brain activity patterns.

In addition to meditation and sauna, music can help us relax and reduce stress [9-11]. The power of music, especially its impact on human emotions and cognition, has long been explored. Existing research has consistently demonstrated that music can not only facilitate relaxation but also induce notable alterations in brain activity [12,13]. However, there is limited empirical evidence on the specific enhancement of theta and alpha frequencies by musical stimuli.

Auditory beat stimulation (ABS) is a non-invasive neuromodulation method that presents a potential solution. It employs auditory stimuli to generate combination tones, encompassing binaural or monaural beats. These beats span across various frequency domains, such as theta (4–8 Hz), alpha (8–13 Hz), and gamma (30–50 Hz), etc. The primary objective of these auditory stimuli is to elicit a neural frequency following response in the listener [14-16]. Monaural beats are generated by superimposing amplitude-modulated signals of closely related frequencies, which can be delivered to a single ear or both ears simultaneously. This phenomenon originates from the interference patterns created when two sound waves with slightly different frequencies are played within a singular auditory channel. The phenomenon is created when two tones with close frequencies are superimposed. This overlapping results in a new frequency, discerned as a rhythmic pulsation, representing the difference in frequency (Hz) between the two original tones [17]. For example, combining tones of 400 Hz and 405 Hz in a single channel results in a 5 Hz monaural beat. A previous study suggested that monaural beat stimulation may help regulating state anxiety and enhancing well-being [18]. However, this approach has its limitations, as such auditory stimulation can be overly monotonous, potentially leading to discomfort or distraction.

Consequently, integrating “normal” music with monaural auditory stimulation might be a promising strategy to brainwave entrainment. This synergy might enhance the overall experience, making it more pleasant and potentially more efficacious in modulating specific brain active patterns, such as the alpha and theta frequency. Thus, this study aimed to devise a novel form of auditory stimulus by combining music with monaural beats, allowing individuals to reach a state of mental and physical well-being similar to going to the sauna. First, we composed a soothing piece of music. Next, after analyzing its frequency spectrum, we incorporated specific pure tones to enhance the brain’s alpha and theta wave activities. Subsequently, we compared the effects of our modified music, regular music, and a silent condition to validate the efficacy of our monaural beatinfused music and determine if it could replicate the benefits derived from a sauna session.

## Materials and Methods

### Participants

Eight healthy participants (4 men, age 21–41 years old) were recruited for participation in the experiment. All participants were right-handed and none had hearing or speech disorders. The experiment consisted of three conditions: modified music (regular music combined with a specific pure tone, henceforth referred to as “music+” condition), regular music (hereafter referred to as “music” condition), and silence. Each participant received the three types of stimuli on different days. To ensure fairness and avoid bias, the order of participation in each condition was balanced among participants.

Informed consent was obtained from all participants prior to the study. All experimental procedures were approved by the Shiba Palace Clinic Ethics Review Committee. Additionally, all procedures were in accordance with the Helsinki Declaration of 1964 and its later amendments

### Procedure

The experimental process included a pre-test phase, a music listening phase (or nothing in the silent condition), and a post-test phase (Fig. 1). Both in the pre-test and post-test phase, we measured scalp electroencephalograms (EEG) and heart rate (HR) during an auditory oddball task. Additionally, we measured salivary alpha-amylase activity (SAA) to assess stress levels using a biosensor (Salivary Amylase Monitor, NIPRO Corporation, Japan) before and after music listening. Further, after the pre-test phase, we recorded 5 min of EEG data under silent conditions as baseline and conducted a questionnaire survey related to physical and mental states after the recording.

**Figure 1:**
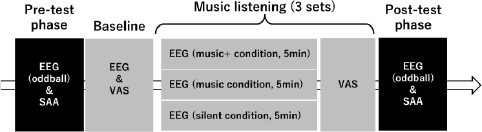
Experimental procedure.

The music listening phase was divided into three parts, each consisting of 5 min of music listening/silence followed by a questionnaire survey. The questionnaire survey consisted of 14 items, extracted from our previous research. These items were chosen because they showed significant changes before and after sauna bathing [8].

### Electroencephalographic recording

In the auditory oddball task, a series of stimuli (total, 200), consisting of standard tones at 1000 Hz (80% of the total) and infrequent “target” tones at 2000 Hz (20% of the total), with a 1.5 s onset of each stimulus were presented. These auditory stimuli, which lasted for 50 ms (with a 5 ms rise/fall time), were delivered through headphones at a sound pressure level of 60 dB. The stimuli were presented pseudo-randomly, ensuring that ≥2 standard stimuli were presented before each “target” stimulus. Participants were asked to press a response button with their preferred hand when detecting the target stimuli. Reaction times (RTs) were measured in ms, and the average of correct responses was calculated. EEG was recorded using a Wireless Biosignal Amplifier System (Polymate Mini AP108; Miyuki Giken Co., Ltd., Tokyo, Japan) and solid gel electrodes (METS INC., Chiba, Japan). We recorded from Cz, T3, and T4 electrodes at a sampling rate of 500 Hz, in accordance with the international 10–20 system. Additionally, ground and reference electrodes were placed on the left and right mastoids, respectively. Electro-ocular (EOG) activity was assessed using an electrode placed at the upper-outer edge of the left eye to measure eye blinks and vertical eye movements. An electrocardiogram was obtained using an electrode placed on the left upper chest to measure the HR. Recorded data were filtered using 0.1 Hz low and 40 Hz high cutoff filters.

During the oddball task, the epoch for the standard or target stimulus began 100 ms before stimulus presentation (baseline) and continued for 600 ms after stimulus onset. Epochs in which the EEG or EOG signal > ±50 μV due to vertical eye movements and muscle contraction artifacts were automatically rejected. Then, standard and target event-related potentials (ERPs) were obtained by averaging the epochs for each stimulus. The difference waveform was calculated by subtracting the standard ERP from the target ERP. We calculated the area of MMN as the negative peak in the difference waveform at 100–250 ms and P300 as the positive peak in the difference waveform at 240–400 ms from stimulus onset.

At baseline and during the music listening/silence period, participants were asked to sit with their eyes closed for 5 min for brain activity measurement. Brain activity data were divided into equally sized segments (10 s) and fast Fourier transformations were conducted. Each participant’s data were averaged across epochs for Cz, T3, and T4 electrodes, respectively, and the gross absolute spectral power for the theta (4–8 Hz) and alpha (8–12 Hz) frequency bands was computed.

### Musical stimulation

One of the authors (KT) composed original music as stimulus for this study using Ableton Live software. The music featured bass, synthesizer, and piano sounds. We utilized sound instruments within the software: “Ambient Encounters Bass” for the bass, “Simple One Pad,” “The Greatest Pad,” and “Megaphatness Pad” for the synthesizers, and “Modular Pianos: 11 Tape Pianos” for the piano. The tones C (Do), C# (Do#), and E (Mi) were played using bass sounds to create low frequency growling sounds. The chord Cadd9, consisting of C (Do), E (Mi), G (Sol), D (Re) tones, was played using synthesizer sounds. The arpeggio melodies consisting of C (Do), D (Re), E (Mi), D (Re) were played with the piano sounds. The results of FFT and frequency analysis of the music envelope are shown in Figure 2. This original music served as stimulus for the music condition.

**Figure 2:**
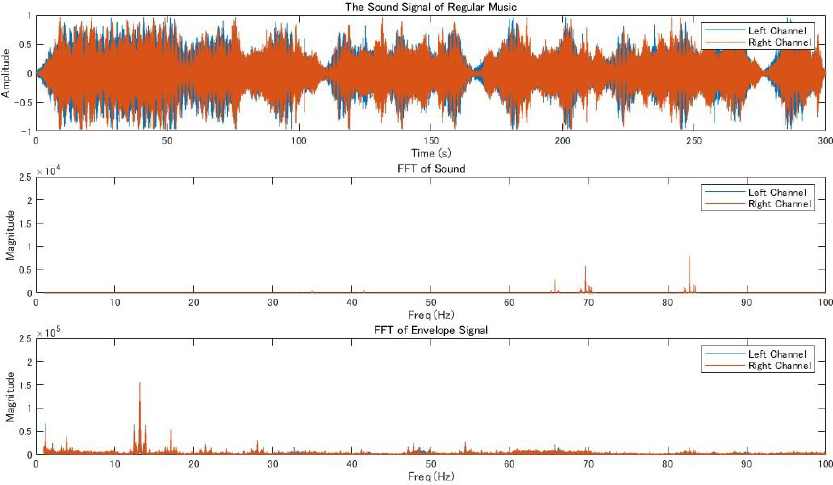
Top, partial waveform of sound stimulus in music condition; middle, frequency response; and bottom, frequency response of the envelope.

Here, we describe our novel auditory stimulus combining original music with pure tones. We define the peak frequency of the original music as *θ*_1_, and that of combining it with a pure tone as *θ*_2_; therefore, we can express the combined sound (S) as

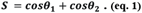

Then, we converted S to an analytical signal by the Hilbert transform and calculated its absolute values to obtain the envelope (E), expressed as

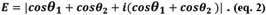

Then, expanding the equation in eq. 2 yields the following equation (ep. 3), which indicates that the peak frequency of E is *θ*_1_ − *θ*_2_ .

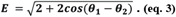

We found peaks around 82 Hz in the Fourier transform of the original music that we created (Fig. 2). Therefore, θ_1_ of the original music was around 82 Hz. To shift the power peak of the music envelope into the theta band (around 7 Hz), we added a 75 Hz pure tone (θ_2_), to the original music. The frequency response and frequency response of the envelope for the modified music is shown in Figure 3. In the frequency response of the envelope, clear peaks in the theta band were found. This music was used as stimulus for the music+ condition.

**Figure 3:**
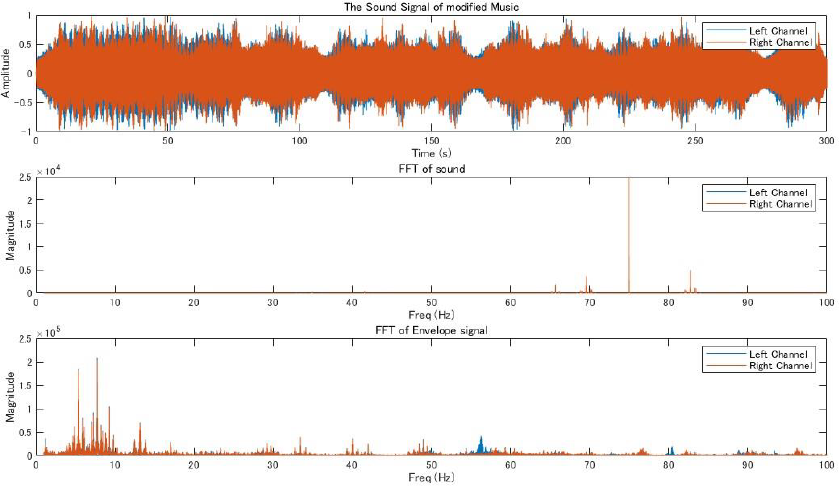
Top, partial waveform of sound stimulus in music+ condition; middle, frequency response; and bottom, frequency response of the envelope.

### Statistical analysis

Ambient For the EEG data recorded during the oddball task in the pre-test and post-test phases, a three-way analysis of variance (ANOVA) was conducted to determine the effects of music listening (pre-test vs. post-test) as a within-subject factor with two levels, conditions (music+ vs. music vs. silent) as a within-subject factor with three levels, and electrode location (Fz vs. T3 vs. T4) as a within-subject factor with three levels.

For the EEG data recorded during the baseline and music listening phases, a three-way ANOVA was conducted to determine the effects of music listening (pre vs. post1 vs. post2 vs. post3) as a within-subject factor with four levels, conditions (music plus vs. music vs. silent) as a within-subject factor with three levels, and electrode location (Fz vs. T3 vs. T4) as a within-subject factor with three levels.

A two-way repeated measures ANOVA was conducted to evaluate changes in participants’ subjective feelings about their physical and mental states. The factors considered were conditions (music plus vs. music vs. silent) and music listening set (pre vs. post1 vs. post2 vs. post3).

A two-way repeated measures ANOVA was conducted to evaluate other effects of music listening. The factors considered were conditions (music plus vs. music vs. silent) and stage (pre-test vs. post-test), which were expected to influence cognitive performance (measured by RT), SAA levels, and HR.

Here, as we aimed to examine the effects of listening to music, we only focused on the main effects of the set and their interactions. If significant effects were identified, the least significant difference method was employed as a post-hoc test for multiple comparisons within each repeated-measure ANOVA. The threshold for significance was set at p < 0.05.

## Results

### Effects of music on cognitive processing and neural oscillation

The We investigated potential changes in cognitive processing after music listening. Utilizing EEG data gathered during the pre-test and post-test phases, we compared the P300 and MMN amplitudes across participants under the three different conditions (Fig. 4).

**Figure 4:**
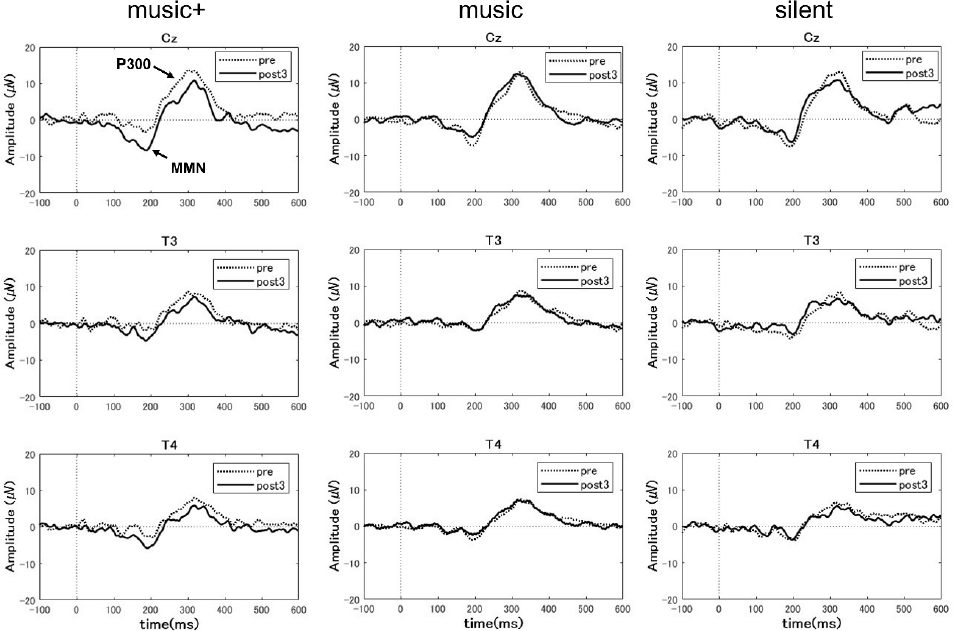
Grand-averaged event-related potential (ERP) difference waveform at three EEG sites (Cz, T3 and T4) for auditory tasks before (pre) and after (post3) music listening across conditions.

For the MMN area, the three-way ANOVA analysis revealed a significant main effect of electrode location (F (2, 28) = 7.94, p < 0.01) and a significant interaction between condition and stage (F (2, 28) = 10.23, p < 0.01). No significant effects were observed for other interactions. The post-hoc tests indicated that, the MMN area significantly increased after listening to music (F (1, 7) = 66.48, p < 0.01) only in the music+ condition. No significant differences were observed in the other conditions (music condition: F (1, 7) = 1.82, n.s; silent condition: F (1, 7) = 3.28, n.s Fig. 5).

**Figure 5:**
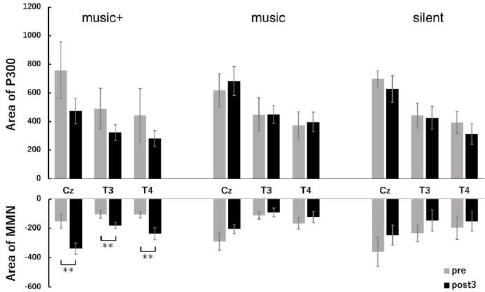
MMN and P3 areas at three locations (Cz, T3, T4) before (pre) and after music listening (post3) across conditions.

A three-way ANOVA with a within-subjects design conducted on P300 area data showed a significant difference only for the electrode location factor (F (2, 28) = 25.77, p < 0.01). No significant differences were detected for other factors or any interactions.

We also evaluated changes in neural oscillations during music listening. Using EEG data gathered at baseline (pre) and throughout each music listening set, we computed the spectral power across the three conditions. Furthermore, we calculated and compared the overall absolute spectral power of the theta and alpha frequency bands for each condition (Fig. 6).

**Figure 6:**
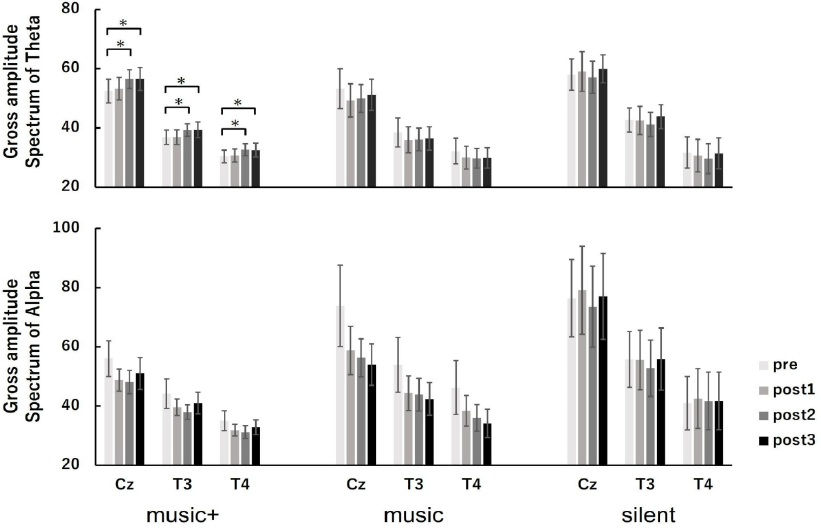
Amplitude spectral power of Theta and Alpha for the Cz, T3 and T4 electrodes at baseline (pre) and at each music listening set (post1, post2, post3) across conditions.

**Figure 7:**
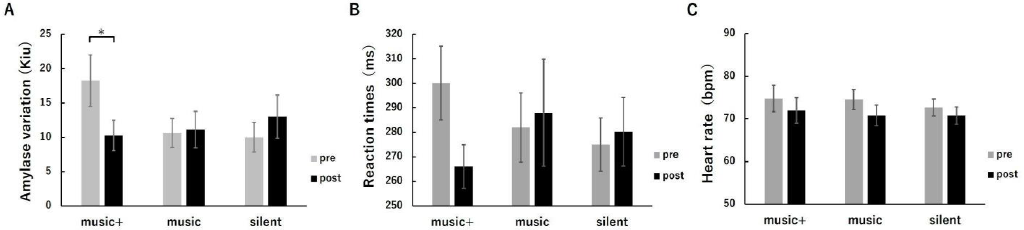
Average SAA (A), RT (B), and HR(C) before (pre) and after music listening (post) across conditions.

Regarding the theta power, the three-way ANOVA revealed a significant main effect for electrode location (F (2, 84) = 133.81, p < 0.01) and a marginally significant interaction between condition and set (F (6, 84) = 1.95, p < 0.1). No significant differences were found for other factors or any interactions. The analysis of condition × set interaction indicated a marginally significant in theta power on set for only the music+ condition (F (3, 21) = 2.65, p < 0.1). However, no significance in theta power on set was observed in either the music (F (3, 21) = 1.37, n.s) or silent conditions (F (3, 21) = 0.68, n.s). A simple main effects test for the set showed a significant increase at post2 and post3 compared with baseline (MSe = 22.3425, p < 0.05) in the music+ condition.

Regarding the alpha power, the three-way ANOVA revealed a significant main effect for set (F (3, 84) = 3.59, p < 0.05) and electrode location (F (2, 84) = 53.20, p < 0.01), a significant interaction between set and electrode location (F (6, 84) = 3.95, p < 0.01) and a marginally significant interaction between condition and electrode location (F (4, 84) = 2.31, p < 0.1). No significant differences were found for condition factors or the interaction between condition and set.

### Effects of music on cognitive performance and physiological parameters

The two-way repeated measures ANOVA for SAA revealed a significant interaction between conditions and stages (F (2, 14) = 4.32, p < 0.05). No significant effects were observed for the group or any interactions. A simple main effects analysis for the stage indicated that the SAA significantly decreased only in the music+ condition (F (1, 7) = 17.40, p < 0.01), while no significant change was observed in the other conditions (music condition: F (1, 7) = 0.03, n.s; silent condition: F (1, 7) = 0.82, n.s). The two-way repeated measures

ANOVA for HR showed no significant interaction between conditions and stages or any significant effects for any factor. The two-way repeated measures ANOVA for RT revealed no significant interaction between conditions and stages. Although a significant effect was observed for stages (F (1, 7) = 10.79, p < 0.05), no significant differences were found among conditions (F (2, 14) = 0.12, n.s).

### Effects of music on subjective feelings

To evaluate the impact of music listening on participants’ subjective perceptions of their physical and mental states, we compared the questionnaire scores at baseline and after each music listening session for all participants across each condition (Fig. 8).

**Figure 8:**
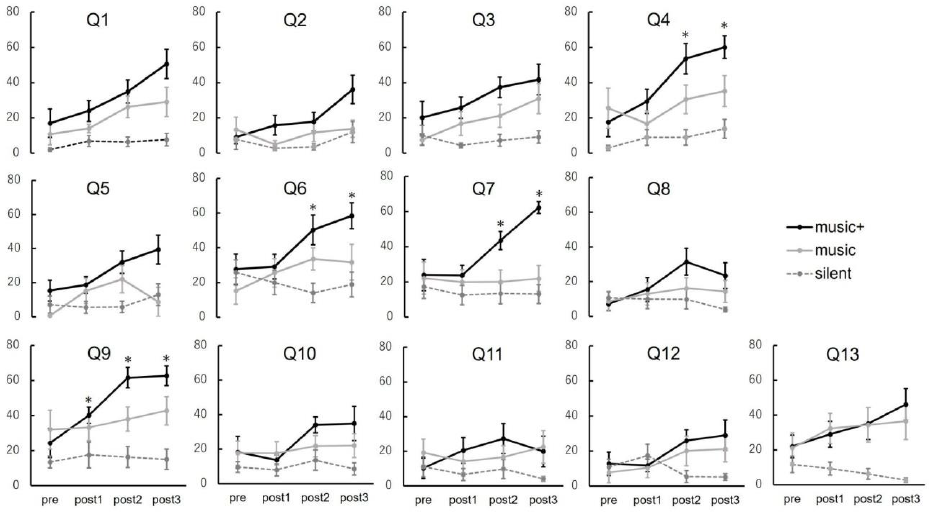
Average scores of questions at baseline (pre) and at each music listening set (post1, post2, post3) across conditions.

The outcomes of the two-way repeated measures ANOVA are shown in Table 1. Out of all 13 questions, significant interactions were observed in the results of four questions. The analysis of condition × stage interaction indicated that the average scores increased significantly only in the music+ condition. Specifically, these four questions are; “Q4: I felt as if I was floating” (F (3,21) = 6.70, p < 0.01), “Q6: I felt isolated from everything and everyone” (F (3,21) = 5.16, p < 0.01), “Q7: my muscles feel loose” (F (3,21) = 8.40, p < 0.01) and “Q9: I’m feeling very relaxed” (F (3,21) = 15.90, p < 0.01). For the questions “My heart is beating faster than usual” and “I have tension in my back,” no significant interactions or main effects were observed. However, for the other questions, there was ≥1 significant main effect either in the set or condition factors.

**Table 1:**
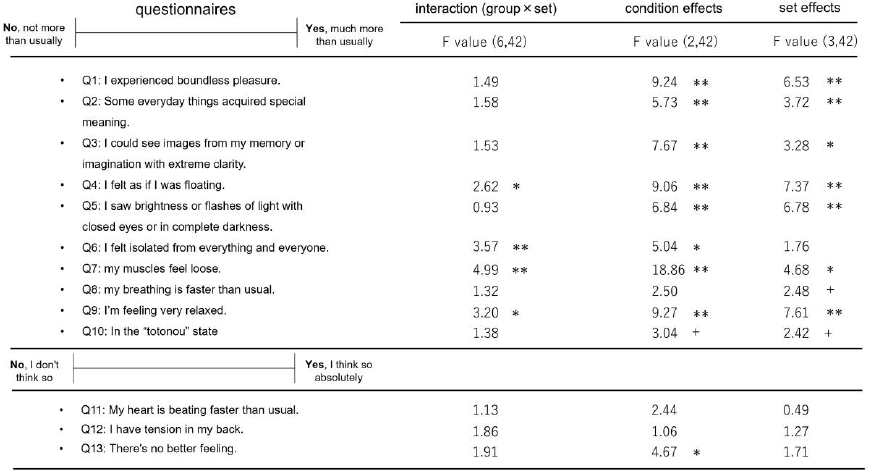
Results of two-way repeated measures ANOVA for all questions in our questionnaires.

## Discussion

In this study, we developed an innovative auditory stimulation technique by merging music with monaural beats, aiming to emulate the therapeutic benefits of a sauna session. This unique blend, which superimposes specific pure tones onto a soothing music, is tailored to amplify the neural activities in the thetaband. By comparing the effects of our modified music, regular music, and silent conditions, we observed that only listening to the modified music significantly increased the MMN amplitude while the SAA levels significantly decreased. Concurrently, only participants exposed to this modified music demonstrated a significant enhancement in theta power. The questionnaire results further highlighted significant differences in the responses of participants under the monaural beat music condition compared to the other conditions.

In this study, the significant increase in MMN amplitude observed during the post-test phase aligns with previous findings from a sauna study [13]. MMN is considered to be attention-dependent; its elicitation can occur regardless of whether participants consciously direct their attention toward the auditory stimulus [19-21]. Thus, MMN is a reliable metric for pre-attentive auditory indices. The increase in MMN amplitude suggests an enhanced sensitivity to auditory stimuli. Additionally, the results from the auditory oddball task indicated a slight reduction in both P300 amplitude and RT after listening to the modified music. This is in contrast to the effects observed after sauna exposure, suggesting that the impact of music listening might not be as pronounced as that of sauna sessions.

In terms of neural oscillations, during music listening of the last two music sets, participants’ theta power significantly increased compared to the baseline, aligning with the findings from a previous sauna study [13]. Theta waves are associated with deep relaxation. Many advanced meditators exhibit increased theta wave activity during their practice, especially as they enter states of profound tranquility [22,23]. This is consistent with the questionnaire results for the statement “I’m feeling very relaxed.” When a person detaches from the external world and immerses in introspection, the brain is likely to enter a theta state. This aligns with the questionnaire results for the statement “I felt isolated from everything and everyone.” It has been suggested that when the brain is in a theta state, it can more easily connect distant ideas, facilitating creative thinking [24]. Additionally, several studies have emphasized the crucial role of theta rhythm in cognitive processes [25]. It is becoming increasingly evident that theta waves are not merely a passive state but actively promote various aspects of cognition. The observed increase in MMN amplitude during the post-test phase might be related to the enhanced theta activity during the music listening process. In our experiments, only the music+ condition increased the power of the theta waves. However, unlike in the sauna research, alpha waves did not increase. This may be because the peak of the envelope of the modified music was around 7 Hz. Since 7 Hz falls within the theta wave band, the theta power should be enhanced, but the alpha power would not. On the other hand, the peak of the envelope for the music in the music condition is around 13 Hz, which would in principle increase alpha waves. However, according to our results, they did not increase but decreased. In contrast to the sauna study, the alpha power did not correlate with any experimental condition, showing a general decline across conditions. Since all experiments were conducted with participants’ eyes closed, participants may easily get drowsy. Some studies have indicated that as individuals transition from a wakeful state to a sleep state, alpha activity tends to decrease [26,27]. Thus, the reduction in alpha power might be attributed to participants gradually transitioning into sleep.

Regarding the questionnaire survey, we selected the questions from the sauna study for which significant interactions were observed. In other words, these questions reflect the subjective effects of the sauna. The present results indicated no significant changes only for the questions “My heart is beating faster than usual” and “I have tension in my back.” For other questions, there was ≥1 significant main effect either in the set or condition factors. This indicates that the sensation of an accelerated HR and tension in the back might be attributed to the alternating hot and cold effects experienced during the sauna sets. Our findings suggest that listening to music is unlikely to replicate these effects.

After completing the experiments under all conditions, we verbally asked the participants if they noticed any anomalies in the two pieces of music they heard. No one noticed any difference between the regular music and the modified music.

Despite the different trends in alpha activity changes between the music+ condition in the present study with the sauna condition in our previous study, there is consistency in the significant increase of theta activity and the significant increase in MMN amplitude. This suggests that, although not fully matching the “totonou” state, we may have successfully created a state somewhat close to it. This fusion strategy presents several advantages. First, it is an accessible relaxation method without the need for specialized settings such as saunas. Furthermore, integrating music with monaural beats did not only make the experience more enjoyable but in the future it could be modified to personal musical preferences.

Nevertheless, our study offers numerous directions for further development and exploration. While our study focused on replicating the benefits derived from a sauna session, future research could investigate its long-term effects and compare them with other relaxation methodologies. Moreover, a broader range of musical genres and styles could be applied to understand their varying impacts on brain activity. Last, to refine and optimize the technique for diverse populations, the individual variability in response to these auditory stimuli warrants further exploration.

In conclusion, our newly developed stimulation protocol for relaxation integrates the demonstrated benefits of music with ABS. As society gravitates toward holistic wellbeing, such innovations hold significant promise in enhancing mental health and overall well-being.

## Supporting information

Supplementary Data: Sound stimuli

## Acknowledgements

We would like to thank Enago (www.enago.jp) for English language editing.

## Author contributions

Conceptualization: Y. N., Y. I., and S. F.

Data curation: C. M.

Formal analysis: C. M.

Funding acquisition: Y. I.

Investigation: C. M., and S. F.

Methodology: C. M., Y. N., Y. I., and S. F.

Project administration: Y. N., Y. I., and S. F.

Resources: C. M., K. T., and S. F.

Software: C. M., K. T., and S.F.

Supervision: Y. N., Y. I., and S. F.

Validation: C. M., and S. F.

Visualization: C. M. and S.F.

Writing – original draft: C. M.

Writing – review & editing: C. M., K. T., Y. N., and S. F.

## Supplementary Data: Sound stimuli

